## Supplementary material for "Single-cell transcriptome sequencing for opening the blood-brain barrier through specific mode electroacupuncture stimulation": Supplementary File 1.docx (from another version of this preprint: version 1, 2025-08-03)

**Supplementary File 1. Pathway analysis for genes downregulated by SMES only in EC_cluster0**

| **Gene Name** | **Gene Functional Annotation** | **Q.value** |
| --- | --- | --- |
| Aif1 | Cellular response to hydroperoxide | 0.016051416 |
|  | Macrophage activation | 0.017010456 |
|  | Inflammatory response | 0.017251057 |
|  | Positive regulation of monocyte chemotaxis | 0.022879878 |
|  | Cellular response to extracellular stimulus | 0.027332483 |
|  | Response to electrical stimulus | 0.035567045 |
|  | Positive regulation of nitric oxide biosynthetic process | 0.037014415 |
|  | Cellular response to hormone stimulus | 0.038907022 |
|  | Actin filament bundle assembly | 0.039704501 |
|  | Positive regulation of T cell proliferation | 0.047704487 |
|  | Phagocytic cup | 0.027332483 |
| Pycard | Positive regulation of chemokine production | 0.00223187 |
|  | Positive regulation of interleukin-6 production | 0.009187487 |
|  | Positive regulation of interleukin-10 production | 0.015064383 |
|  | Cellular response to lipopolysaccharide | 0.015295529 |
|  | Regulation of tumor necrosis factor-mediated signaling pathway | 0.015295529 |
|  | Positive regulation of cysteine-type endopeptidase activity | 0.015295529 |
|  | Interleukin-1 beta production | 0.016051416 |
|  | Macropinocytosis | 0.016051416 |
|  | Inflammatory response | 0.017251057 |
|  | Negative regulation of interferon-beta production | 0.017251057 |
|  | Activation of cysteine-type endopeptidase activity | 0.017251057 |
|  | Positive regulation of interleukin-8 production | 0.017251057 |
|  | Positive regulation of T cell activation | 0.022879878 |
|  | Positive regulation of activated T cell proliferation | 0.024146921 |
|  | Positive regulation of interleukin-1 beta production | 0.028164725 |
|  | Positive regulation of interleukin-6 production | 0.02896625 |
|  | Negative regulation of canonical NF-kappab signal transduction | 0.03839198 |
|  | Positive regulation of type II interferon production | 0.040489387 |
|  | Tumor necrosis factor-mediated signaling pathway | 0.041261961 |
|  | Regulation of inflammatory response | 0.047014148 |
|  | Positive regulation of tumor necrosis factor production | 0.047704487 |
|  | Positive regulation of cysteine-type endopeptidase activity involved in apoptotic process | 0.047704487 |
|  | Cytosol | 0.034959432 |
|  | NLRP1 inflammasome complex | 0.014691841 |
|  | AIM2 inflammasome complex | 0.015295529 |
|  | NLRP3 inflammasome complex | 0.016051416 |
|  | Ikappab kinase complex | 0.017143782 |
|  | Interleukin-6 receptor binding | 0.016051416 |
|  | BMP receptor binding | 0.016051416 |
|  | Cysteine-type endopeptidase activity involved in apoptotic process | 0.017251057 |
| Il33 | Positive regulation of chemokine production | 0.00223187 |
|  | Positive regulation of interleukin-6 production | 0.009187487 |
|  | Microglial cell proliferation | 0.014691841 |
|  | Microglial cell activation involved in immune response | 0.014691841 |
|  | Positive regulation of interleukin-13 production | 0.016051416 |
|  | Positive regulation of interleukin-5 production | 0.016051416 |
|  | Macrophage activation involved in immune response | 0.017251057 |
|  | Positive regulation of interleukin-4 production | 0.019540379 |
|  | Positive regulation of macrophage activation | 0.020702823 |
|  | Negative regulation of type II interferon production | 0.025130843 |
|  | Positive regulation of cytokine production | 0.027332483 |
|  | Positive regulation of inflammatory response | 0.047704487 |
|  | Cytosol | 0.034959432 |
|  | Transport vesicle | 0.045940999 |
| Ltc4s | Cellular response to lipopolysaccharide | 0.015295529 |
|  | Leukotriene biosynthetic process | 0.016051416 |
|  | Leukotriene metabolic process | 0.017010456 |
|  | Response to lipopolysaccharide | 0.017143782 |
|  | Nuclear outer membrane | 0.019540379 |
|  | Leukotriene-C4 synthase activity | 0.015295529 |
|  | Glutathione binding | 0.020702823 |
|  | Glutathione peroxidase activity | 0.026074999 |
|  | Glutathione transferase activity | 0.037860086 |
