## Supplementary material for "Single-cell transcriptome sequencing for opening the blood-brain barrier through specific mode electroacupuncture stimulation": Supplementary File 2.docx (from another version of this preprint: version 1, 2025-08-03)

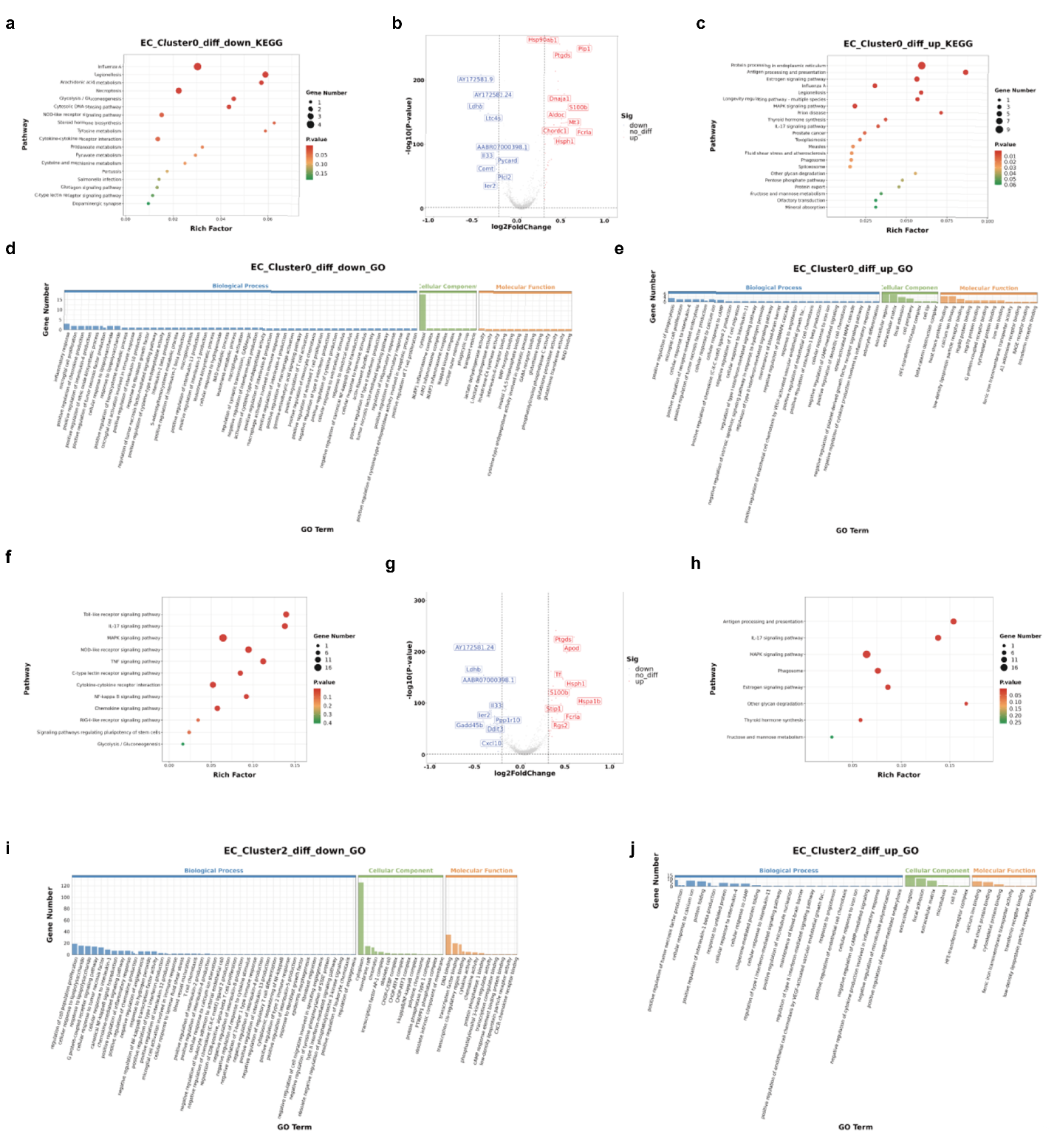

**Supplementary File 2.** **Differential gene and enrichment analysis results of EC_Cluster0 and EC_Cluster2, related to Figure 7. (A)** KEGG enrichment results for down-regulated genes in EC_Cluster0. **(B)** EC_Cluster0 differential gene volcano map. **(C)** KEGG results of EC_Cluster0 upregulated genes. **(D)** GO results of EC_Cluster 0 downregulated genes. **(E)** GO results of EC_Cluster0 upregulated genes. **(F)** KEGG results of EC_Cluster2 downregulated genes. **(G)** EC_Cluster2 differential gene volcano map. **(H)** KEGG results of EC_Cluster2 upregulated genes. **(I)** GO results of EC_Cluster2 downregulated genes. **(J)** GO results of EC_Cluster2 upregulated genes. GO, Gene Ontology; KEGG, Kyoto Encyclopedia of Genes and Genomes
