## Supplementary material for "Single-cell transcriptome sequencing for opening the blood-brain barrier through specific mode electroacupuncture stimulation": Supplementary File 4.docx (from another version of this preprint: version 1, 2025-08-03)

**Supplementary File 4. Pathway analysis for genes upregulated by SMES only in EC_cluster0**

| **Gene Name** | **Gene Functional Annotation** | **Q.value** |
| --- | --- | --- |
| S100b | astrocyte differentiation | 0.047090205 |
|  | calcium ion binding | 0.036073527 |
|  | RAGE receptor binding | 0.03999733 |
| Dnaja1 | heat shock protein binding | 4.6715E-05 |
|  | low-density lipoprotein particle receptor binding | 0.000203237 |
|  | Hsp70 protein binding | 0.02244097 |
|  | G protein-coupled receptor binding | 0.02250692 |
| Aldoc | cytoskeletal protein binding | 0.033981815 |
| Hsph1 | negative regulation of intrinsic apoptotic signaling pathway in response to hydrogen peroxide | 0.027868317 |
|  | negative regulation of p38MAPK cascade | 0.034043282 |
|  | heat shock protein binding | 4.6715E-05 |
| Hsp90ab1 | cellular response to interleukin-4 | 0.008538995 |
|  | regulation of type I interferon-mediated signaling pathway | 0.027868317 |
|  | regulation of type II interferon-mediated signaling pathway | 0.034043282 |
| Stip1 | Hsp90 protein binding | 0.01704401 |
|  | Hsp70 protein binding | 0.02244097 |
| Hspa5 | cellular response to interleukin-4 | 0.008538995 |
|  | cellular response to calcium ion | 0.024673147 |
|  | cellular response to cAMP | 0.027391704 |
|  | extracellular matrix | 0.00015723 |
|  | focal adhesion | 0.027868317 |
|  | heat shock protein binding | 4.6715E-05 |
| Hspa8 | positive regulation of phagocytosis | 0.017880273 |
|  | extracellular matrix | 0.00015723 |
|  | heat shock protein binding | 4.6715E-05 |
|  | G protein-coupled receptor binding | 0.02250692 |
|  | A1 adenosine receptor binding | 0.03999733 |
| P4ha1 | iron ion binding | 0.049215868 |
| Calr | positive regulation of phagocytosis | 0.017880273 |
|  | positive regulation of dendritic cell chemotaxis | 0.043359962 |
|  | extracellular matrix | 0.00015723 |
|  | focal adhesion | 0.027868317 |
|  | calcium ion binding | 0.036073527 |
|  | iron ion binding | 0.049215868 |
| Tf | positive regulation of receptor-mediated endocytosis | 0.009996979 |
|  | cellular response to cAMP | 0.027391704 |
|  | cellular response to iron ion | 0.043359962 |
|  | extracellular region | 0.002111889 |
|  | HFE-transferrin receptor complex | 0.036073527 |
|  | cell tip | 0.036073527 |
|  | ferric iron transmembrane transporter activity | 0.015409729 |
|  | transferrin receptor binding | 0.047090205 |
| Clu | positive regulation of receptor-mediated endocytosis | 0.009996979 |
|  | microglial cell proliferation | 0.02250692 |
|  | extracellular matrix | 0.00015723 |
|  | extracellular region | 0.002111889 |
|  | cell periphery | 0.002490811 |
|  | low-density lipoprotein particle receptor binding | 0.000203237 |
| Cryab | stress-activated MAPK cascade | 0.047090205 |
|  | cytoskeletal protein binding | 0.033981815 |
| Mbp | positive regulation of chemokine (C-X-C motif) ligand 2 production | 0.015409729 |
|  | maintenance of blood-brain barrier | 0.034043282 |
|  | cell periphery | 0.002490811 |
| Hspb1 | cellular response to interleukin-11 | 0.02250692 |
|  | response to angiotensin | 0.034043282 |
|  | positive regulation of endothelial cell chemotaxis by VEGF-activated vascular endothelial growth factor receptor signaling pathway | 0.036073527 |
|  | positive regulation of endothelial cell chemotaxis | 0.036073527 |
|  | positive regulation of interleukin-1 beta production | 0.03999733 |
|  | positive regulation of tumor necrosis factor production | 0.043359962 |
|  | extracellular matrix | 0.00015723 |
|  | focal adhesion | 0.027868317 |
| Lyve1 | cell periphery | 0.002490811 |
| Cxcl14 | extracellular region | 0.002111889 |
