## Supplementary material for "Single-cell transcriptome sequencing for opening the blood-brain barrier through specific mode electroacupuncture stimulation": Supplementary File 6.docx (from another version of this preprint: version 1, 2025-08-03)

**Supplementary File 6. Pathway analysis for genes upregulated only in EC_cluster2**

| Psap | Extracellular matrix | 0.00018412 |
| --- | --- | --- |
|  | Extracellular region | 0.000548325 |
| Mt-co3 | Oxidoreduction-driven active transmembrane transporter activity | 0.014827033 |
| Pdia4 | Chaperone-mediated protein folding | 2.31652E-07 |
|  | Protein folding | 1.98193E-05 |
