## Supplementary material for "Single-cell transcriptome sequencing for opening the blood-brain barrier through specific mode electroacupuncture stimulation": Supplementary File 7.docx (from another version of this preprint: version 1, 2025-08-03)

**Supplementary File 7. Pathway analysis for genes downregulated only in EC_cluster2**

| Nfkbia | regulation of cell population proliferation | 0.003486927 |
| --- | --- | --- |
|  | canonical NF-kappaB signal transduction | 0.017096451 |
|  | positive regulation of inflammatory response | 0.020796191 |
|  | response to lipopolysaccharide | 0.023336962 |
|  | negative regulation of NF-kappaB transcription factor activity | 0.023896228 |
|  | cytoplasmic sequestering of NF-kappaB | 0.029332997 |
|  | cellular response to tumor necrosis factor | 0.035477619 |
|  | cytoplasm | 0.017096451 |
|  | I-kappaB/NF-kappaB complex | 0.035477619 |
| Klf4 | regulation of cell population proliferation | 0.003486927 |
|  | negative regulation of leukocyte adhesion to arterial endothelial cell | 0.017096451 |
|  | negative regulation of chemokine (C-X-C motif) ligand 2 production | 0.017096451 |
|  | negative regulation of interleukin-8 production | 0.020796191 |
|  | negative regulation of angiogenesis | 0.022740408 |
|  | negative regulation of NF-kappaB transcription factor activity | 0.023896228 |
|  | negative regulation of response to cytokine stimulus | 0.023896228 |
|  | cellular response to laminar fluid shear stress | 0.026777783 |
|  | epidermis morphogenesis | 0.029332997 |
|  | negative regulation of cell migration involved in sprouting angiogenesis | 0.035477619 |
|  | obsolete negative regulation of phosphatidylinositol 3-kinase signaling | 0.048523805 |
|  | cytoplasm | 0.017096451 |
|  | chromatin | 0.022740408 |
|  | chromatin | 0.036900462 |
|  | phosphatidylinositol 3-kinase regulator activity | 0.029332997 |
| Cxcl2 | regulation of cell population proliferation | 0.003486927 |
|  | positive regulation of leukocyte chemotaxis | 0.012823597 |
|  | chemokine-mediated signaling pathway | 0.017096451 |
|  | response to lipopolysaccharide | 0.023336962 |
|  | cellular response to interleukin-1 | 0.025889974 |
|  | G protein-coupled receptor signaling pathway | 0.026777783 |
|  | cellular response to lipopolysaccharide | 0.045991692 |
|  | obsolete cell | 0.026777783 |
|  | CXCR chemokine receptor binding | 0.012823597 |
|  | chemokine activity | 0.017096451 |
| Id1 | regulation of angiogenesis | 0.017096451 |
|  | regulation of vasculature development | 0.017096451 |
|  | negative regulation of endothelial cell differentiation | 0.035477619 |
|  | endothelial cell morphogenesis | 0.039382949 |
|  | cytoplasm | 0.017096451 |
| Olr1 | response to hydrogen peroxide | 0.023896228 |
|  | membrane raft | 0.023896228 |
| Irf1 | regulation of cell population proliferation | 0.003486927 |
|  | canonical NF-kappaB signal transduction | 0.017096451 |
|  | regulation of CD8-positive, alpha-beta T cell proliferation | 0.017096451 |
|  | cellular response to interleukin-1 | 0.025889974 |
|  | negative regulation of regulatory T cell differentiation | 0.026777783 |
|  | positive regulation of interleukin-12 production | 0.029332997 |
|  | type II interferon-mediated signaling pathway | 0.034978747 |
|  | cellular response to tumor necrosis factor | 0.035477619 |
|  | negative regulation of tyrosine phosphorylation of STAT protein | 0.041674461 |
|  | positive regulation of type I interferon production | 0.048523805 |
|  | cytoplasm | 0.017096451 |
|  | chromatin | 0.022740408 |
| Ddit3 | response to hydrogen peroxide | 0.023896228 |
|  | blood vessel maturation | 0.035477619 |
|  | cytoplasm | 0.017096451 |
|  | CHOP-C/EBP complex | 0.020796191 |
|  | CHOP-ATF4 complex | 0.020796191 |
|  | CHOP-ATF3 complex | 0.020796191 |
|  | transcription factor AP-1 complex | 0.029332997 |
|  | cAMP response element binding protein binding | 0.036900462 |
| Cxcl10 | regulation of cell population proliferation | 0.003486927 |
|  | positive regulation of leukocyte chemotaxis | 0.012823597 |
|  | chemokine-mediated signaling pathway | 0.017096451 |
|  | regulation of T cell chemotaxis | 0.020796191 |
|  | negative regulation of angiogenesis | 0.022740408 |
|  | response to lipopolysaccharide | 0.023336962 |
|  | regulation of endothelial tube morphogenesis | 0.023896228 |
|  | G protein-coupled receptor signaling pathway | 0.026777783 |
|  | T cell chemotaxis | 0.03273337 |
|  | endothelial cell activation | 0.036900462 |
|  | positive regulation of T cell migration | 0.041674461 |
|  | cellular response to lipopolysaccharide | 0.045991692 |
|  | CXCR chemokine receptor binding | 0.012823597 |
|  | chemokine activity | 0.017096451 |
|  | cytokine activity | 0.026777783 |
|  | CXCR3 chemokine receptor binding | 0.026777783 |
| Gadd45b | cytoplasm | 0.017096451 |
| Sertad1 | cytoplasm | 0.017096451 |
| Ppp1r10 | chromatin | 0.022740408 |
|  | chromatin | 0.036900462 |
|  | PTW/PP1 phosphatase complex | 0.036900462 |
| Zfand5 | fibroblast migration | 0.035477619 |
|  | cytoplasm | 0.017096451 |
| H3f3a | polytene chromosome | 0.020796191 |
| Rilpl2 | cytoplasm | 0.017096451 |
|  | obsolete cell | 0.026777783 |
| Rgs16 | G protein-coupled receptor signaling pathway | 0.026777783 |
|  | cytoplasm | 0.017096451 |
|  | membrane raft | 0.023896228 |
|  | obsolete intrinsic component of membrane | 0.045991692 |
| Per3 | cytoplasm | 0.017096451 |
| Cd83 | negative regulation of interleukin-4 production | 0.020796191 |
|  | positive regulation of interleukin-2 production | 0.045991692 |
| Idi1 | cytoplasm | 0.017096451 |
| Tsc22d3 | cytoplasm | 0.017096451 |
