## Supplementary material for "Single-cell transcriptome sequencing for opening the blood-brain barrier through specific mode electroacupuncture stimulation": Supplementary File 8.docx (from another version of this preprint: version 1, 2025-08-03)

**Supplementary File 8. Pathway analysis for genes downregulated only in EC_cluster4**

| Cd74 | positive regulation of ERK1 and ERK2 cascade | 0.001856584 |
| --- | --- | --- |
|  | macrophage migration inhibitory factor signaling pathway | 0.01311054 |
|  | positive regulation of chemokine (C-X-C motif) ligand 2 production | 0.02574415 |
|  | positive regulation of type 2 immune response | 0.027247177 |
|  | positive regulation of monocyte differentiation | 0.030717148 |
|  | membrane | 0.000510691 |
|  | NOS2-CD74 complex | 0.01311054 |
|  | macrophage migration inhibitory factor receptor complex | 0.022461526 |
|  | MHC class II protein complex | 0.028691751 |
|  | macrophage migration inhibitory factor binding | 0.022461526 |
|  | MHC class II protein binding | 0.02574415 |
|  | CD4 receptor binding | 0.030144772 |
|  | nitric-oxide synthase binding | 0.035835211 |
|  | MHC class II protein complex binding | 0.037879909 |
|  | cytokine binding | 0.039302447 |
| RT1-CE10 | antigen processing and presentation of peptide antigen via MHC class I | 2.0785E-06 |
|  | immune response | 0.001787329 |
|  | MHC class I protein complex | 1.14188E-06 |
|  | membrane | 0.000510691 |
| RT1-T24-3 | antigen processing and presentation of peptide antigen via MHC class I | 2.0785E-06 |
|  | immune response | 0.001787329 |
|  | MHC class I protein complex | 1.14188E-06 |
|  | membrane | 0.000510691 |
| RT1-A2 | antigen processing and presentation of peptide antigen via MHC class I | 2.0785E-06 |
|  | immune response | 0.001787329 |
|  | MHC class I protein complex | 1.14188E-06 |
|  | membrane | 0.000510691 |
| Fcgrt | membrane | 0.000510691 |
| RT1-CE4 | antigen processing and presentation of peptide antigen via MHC class I | 2.0785E-06 |
|  | immune response | 0.001787329 |
|  | MHC class I protein complex | 1.14188E-06 |
|  | membrane | 0.000510691 |
| Trem2 | positive regulation of antigen processing and presentation of peptide antigen via MHC class II | 0.000119743 |
|  | phagocytosis, engulfment | 0.000281174 |
|  | positive regulation of ERK1 and ERK2 cascade | 0.001856584 |
|  | positive regulation of C-C chemokine receptor CCR7 signaling pathway | 0.01311054 |
|  | positive regulation of CD40 signaling pathway | 0.01311054 |
|  | dendritic cell differentiation | 0.030717148 |
|  | positive regulation of calcium-mediated signaling | 0.039302447 |
|  | membrane | 0.000510691 |
|  | lipopolysaccharide binding | 0.039302447 |
| Lgals3bp | receptor-mediated endocytosis | 0.025074728 |
| Cldn5 | calcium-independent cell-cell adhesion via plasma membrane cell-adhesion molecules | 0.025074728 |
|  | positive regulation of establishment of endothelial barrier | 0.02574415 |
|  | cell-cell junction assembly | 0.04609774 |
|  | membrane | 0.000510691 |
|  | Schmidt-Lanterman incisure | 0.037808896 |
| Ifitm3 | receptor-mediated endocytosis | 0.025074728 |
|  | response to interferon-alpha | 0.03231788 |
|  | response to interferon-beta | 0.034014558 |
|  | type I interferon-mediated signaling pathway | 0.035835211 |
|  | membrane | 0.000510691 |
| C1qa | complement activation, classical pathway | 0.005686291 |
|  | response to iron ion | 0.047687967 |
| Rtp4 | membrane | 0.000510691 |
