## Supplementary material for "Single-cell transcriptome sequencing for opening the blood-brain barrier through specific mode electroacupuncture stimulation": Supplementary File 9.docx (from another version of this preprint: version 1, 2025-08-03)

**Supplementary File 9. Pathway analysis for genes upregulated only in EC_cluster4**

| Zfp36 | MAPK cascade | 0.00250583 |
| --- | --- | --- |
|  | phosphatidylinositol 3-kinase/protein kinase B signal transduction | 0.002647933 |
|  | cellular response to tumor necrosis factor | 0.003156035 |
|  | p38MAPK cascade | 0.003670272 |
|  | obsolete phosphatidylinositol 3-kinase signaling | 0.012327772 |
|  | cellular response to glucocorticoid stimulus | 0.018962001 |
|  | ERK1 and ERK2 cascade | 0.018962001 |
|  | cellular response to fibroblast growth factor stimulus | 0.019159141 |
|  | cellular response to epidermal growth factor stimulus | 0.026707055 |
|  | negative regulation of interleukin-2 production | 0.032454936 |
|  | vasculogenesis | 0.034405213 |
|  | cellular response to cAMP | 0.034405213 |
|  | cell population proliferation | 0.04027658 |
|  | heat shock protein binding | 1.46638E-08 |
|  | C-C chemokine binding | 0.002647933 |
| Hsp90aa1 | positive regulation of nitric oxide biosynthetic process | 0.021649389 |
|  | extracellular matrix | 0.003517823 |
|  | membrane | 0.026630135 |
|  | nitric-oxide synthase regulator activity | 0.032454936 |
| Cited2 | positive regulation of cell-cell adhesion | 0.009649989 |
|  | vasculogenesis | 0.034405213 |
| Itgb2 | neutrophil chemotaxis | 0.003517823 |
|  | positive regulation of nitric oxide biosynthetic process | 0.021649389 |
|  | membrane raft | 0.004504895 |
|  | membrane | 0.026630135 |
|  | heat shock protein binding | 1.46638E-08 |
|  | ICAM-3 receptor activity | 0.026630135 |
| Tlr3 | positive regulation of inflammatory response | 2.64486E-05 |
|  | positive regulation of interleukin-6 production | 0.003125083 |
|  | positive regulation of interleukin-12 production | 0.01041014 |
|  | cellular response to type II interferon | 0.013764808 |
|  | positive regulation of type III interferon production | 0.018962001 |
|  | microglial cell activation involved in immune response | 0.032454936 |
|  | MyD88-independent toll-like receptor signaling pathway | 0.035660924 |
|  | positive regulation of toll-like receptor signaling pathway | 0.035660924 |
|  | positive regulation of interferon-alpha production | 0.035660924 |
|  | positive regulation of chemokine production | 0.043224513 |
|  | positive regulation of interferon-beta production | 0.047220479 |
| Ptges3 | glucocorticoid receptor signaling pathway | 0.039296238 |
|  | cell population proliferation | 0.04027658 |
| Hexb | astrocyte cell migration | 0.003125083 |
|  | glycosaminoglycan metabolic process | 0.039296238 |
|  | intracellular calcium ion homeostasis | 0.039721568 |
|  | membrane | 0.026630135 |
|  | extracellular space | 0.043224513 |
| Id2 | Peyer's patch development | 0.043224513 |
| Dynll1 | microtubule | 0.019159141 |
|  | membrane | 0.026630135 |
| Igf1 | phosphatidylinositol 3-kinase/protein kinase B signal transduction | 0.002647933 |
|  | obsolete phosphatidylinositol 3-kinase signaling | 0.012327772 |
|  | ERK1 and ERK2 cascade | 0.018962001 |
|  | positive regulation of cell migration | 0.035660924 |
|  | regulation of nitric oxide biosynthetic process | 0.035660924 |
|  | positive regulation of phosphatidylinositol 3-kinase/protein kinase B signal transduction | 0.039721568 |
|  | cell population proliferation | 0.04027658 |
|  | positive regulation of glycolytic process | 0.049977366 |
|  | response to epidermal growth factor | 0.049977366 |
|  | positive regulation of calcineurin-NFAT signaling cascade | 0.049977366 |
|  | extracellular space | 0.043224513 |
| Ppt1 | positive regulation of pinocytosis | 0.032454936 |
|  | pinocytosis | 0.035660924 |
|  | membrane raft | 0.004504895 |
|  | membrane | 0.026630135 |
|  | extracellular space | 0.043224513 |
| St13 | heat shock protein binding | 1.46638E-08 |
|  | Hsp70 protein binding | 0.003156035 |
| Ccr5 | positive regulation of inflammatory response | 2.64486E-05 |
|  | MAPK cascade | 0.00250583 |
|  | release of sequestered calcium ion into cytosol by sarcoplasmic reticulum | 0.002647933 |
|  | chemokine-mediated signaling pathway | 0.002988252 |
|  | positive regulation of interleukin-6 production | 0.003125083 |
|  | positive regulation of cell-cell adhesion | 0.009649989 |
|  | positive regulation of interleukin-1 beta production | 0.017952523 |
|  | G protein-coupled receptor signaling pathway | 0.021046863 |
|  | calcium-mediated signaling | 0.022698002 |
|  | calcium ion transport | 0.040380395 |
|  | positive regulation of cytosolic calcium ion concentration | 0.047220479 |
|  | C-C chemokine binding | 0.002647933 |
|  | chemokine (C-C motif) ligand 5 binding | 0.026630135 |
|  | C-C chemokine receptor activity | 0.049977366 |
