## Supplementary material for "Single-cell transcriptome sequencing for opening the blood-brain barrier through specific mode electroacupuncture stimulation": Supplementary File 10.docx (from another version of this preprint: version 1, 2025-08-03)

**Supplementary File 10. Pathway analysis for genes downregulated only in EC_cluster5**

| **Gene Functional Annotation** | **Q.value** | **Gene Number** |
| --- | --- | --- |
| structural constituent of ribosome | 1.13363E-48 | 49 |
| translation | 4.72863E-39 | 42 |
| cytosolic large ribosomal subunit | 2.15156E-25 | 26 |
| RNA binding | 0.013829115 | 20 |
| cytosolic small ribosomal subunit | 2.67958E-17 | 18 |
| focal adhesion | 5.75812E-09 | 18 |
| cytoplasmic translation | 1.18745E-09 | 11 |
| ribosomal large subunit assembly | 8.48666E-07 | 7 |
| rRNA binding | 1.1968E-05 | 7 |
| rRNA processing | 0.005707569 | 6 |
| mRNA binding | 0.031255076 | 6 |
| ribosomal small subunit assembly | 0.000110129 | 5 |
| 5S rRNA binding | 2.59746E-06 | 5 |
| positive regulation of translation | 0.037400694 | 4 |
| polysome | 0.014750505 | 4 |
| cytoplasmic ribonucleoprotein granule | 0.035284417 | 4 |
| SRP-dependent co-translational protein targeting to membrane | 0.035499568 | 3 |
| small ribosomal subunit | 0.007701374 | 3 |
| large ribosomal subunit | 0.009016838 | 3 |
| small ribosomal subunit rRNA binding | 0.002870644 | 3 |
| translation elongation factor activity | 0.014750505 | 3 |
| positive regulation of cyclic-nucleotide phosphodiesterase activity | 0.002986019 | 2 |
| positive regulation of mitochondrial depolarization | 0.042970815 | 2 |
| Rho GDP-dissociation inhibitor activity | 0.002986019 | 2 |
