## Supplementary material for "Single-cell transcriptome sequencing for opening the blood-brain barrier through specific mode electroacupuncture stimulation": Supplementary File 11.docx (from another version of this preprint: version 1, 2025-08-03)

**Supplementary File 11. Pathway analysis for genes upregulated only in EC_cluster5**

| **Gene Functional Annotation** | **Q.value** | **Gene Number** |
| --- | --- | --- |
| extracellular space | 0.016822971 | 10 |
| G protein-coupled receptor signaling pathway | 0.009472202 | 6 |
| myelin sheath | 0.020927137 | 5 |
| membrane raft | 0.030105854 | 5 |
| leukocyte chemotaxis | 8.90296E-05 | 4 |
| mitochondrial respiratory chain complex I | 0.003917673 | 4 |
| chemokine activity | 0.000281694 | 4 |
| lymphocyte chemotaxis | 0.00149395 | 3 |
| chemokine-mediated signaling pathway | 0.010612598 | 3 |
| neutrophil chemotaxis | 0.016822971 | 3 |
| positive regulation of tumor necrosis factor production | 0.024856709 | 3 |
| regulation of chemotaxis | 0.004761556 | 2 |
| macrophage chemotaxis | 0.026130337 | 2 |
| sprouting angiogenesis | 0.030105854 | 2 |
| positive regulation of interleukin-10 production | 0.036311293 | 2 |
| regulation of cytokine production | 0.036311293 | 2 |
| mitochondrial proton-transporting ATP synthase complex | 0.039574911 | 2 |
| NADH dehydrogenase activity | 0.026130337 | 2 |
| G-protein alpha-subunit binding | 0.036311293 | 2 |
| ionotropic glutamate receptor binding | 0.036311293 | 2 |
| lamellipodium assembly involved in ameboidal cell migration | 0.036311293 | 1 |
| regulation of establishment of blood-brain barrier | 0.036311293 | 1 |
| host cell cytoplasm | 0.036311293 | 1 |
| sodium-dependent organic anion transmembrane transporter activity | 0.036311293 | 1 |
