## Supplementary material for "Single-cell transcriptome sequencing for opening the blood-brain barrier through specific mode electroacupuncture stimulation": Supplementary File 15.docx (from another version of this preprint: version 1, 2025-08-03)

**Supplementary File 15. GO analysis for MG_cluster1 top genes only (S≥2)**

| **GO_ID** | **GO_Term** | **S** |
| --- | --- | --- |
| [GO:0016021](http://amigo.geneontology.org/amigo/term/GO:0016021) | membrane | 5 |
| [GO:0007186](http://amigo.geneontology.org/amigo/term/GO:0007186) | G protein-coupled receptor signaling pathway | 4 |
| [GO:0005615](http://amigo.geneontology.org/amigo/term/GO:0005615) | extracellular space | 3 |
| [GO:0005887](http://amigo.geneontology.org/amigo/term/GO:0005887) | plasma membrane | 3 |
| [GO:0043231](http://amigo.geneontology.org/amigo/term/GO:0043231) | intracellular membrane-bounded organelle | 3 |
| [GO:0045028](http://amigo.geneontology.org/amigo/term/GO:0045028) | G protein-coupled purinergic nucleotide receptor activity | 3 |
| [GO:0009897](http://amigo.geneontology.org/amigo/term/GO:0009897) | external side of plasma membrane | 2 |
| [GO:0019221](http://amigo.geneontology.org/amigo/term/GO:0019221) | cytokine-mediated signaling pathway | 2 |
| [GO:0032809](http://amigo.geneontology.org/amigo/term/GO:0032809) | neuronal cell body membrane | 2 |
| [GO:0035589](http://amigo.geneontology.org/amigo/term/GO:0035589) | G protein-coupled purinergic nucleotide receptor signaling pathway | 2 |
| [GO:0048246](http://amigo.geneontology.org/amigo/term/GO:0048246) | macrophage chemotaxis | 2 |
| [GO:0071407](http://amigo.geneontology.org/amigo/term/GO:0071407) | cellular response to organic cyclic compound | 2 |
