## Supplementary material for "Single-cell transcriptome sequencing for opening the blood-brain barrier through specific mode electroacupuncture stimulation": Supplementary File 16.docx (from another version of this preprint: version 1, 2025-08-03)

**Supplementary File 16. KEGG analysis for MG_cluster1 top genes only (counts top 20)**

| **Pathway_ID** | **Pathway_Name** | **S** |
| --- | --- | --- |
| [rno04060](https://www.kegg.jp/entry/rno04060) | Cytokine-cytokine receptor interaction | 2 |
| [rno00590](https://www.kegg.jp/entry/rno00590) | Arachidonic acid metabolism | 1 |
| [rno04512](https://www.kegg.jp/entry/rno04512) | ECM-receptor interaction | 1 |
| [rno04550](https://www.kegg.jp/entry/rno04550) | Signaling pathways regulating pluripotency of stem cells | 1 |
| [rno04611](https://www.kegg.jp/entry/rno04611) | Platelet activation | 1 |
| [rno04630](https://www.kegg.jp/entry/rno04630) | JAK-STAT signaling pathway | 1 |
