## Supplementary material for "Single-cell transcriptome sequencing for opening the blood-brain barrier through specific mode electroacupuncture stimulation": Supplementary File 18.docx (from another version of this preprint: version 1, 2025-08-03)

**Supplementary File 18. GO analysis for MG_cluster0 top genes only (S≥2)**

| **GO_ID** | **GO_Term** | **S** |
| --- | --- | --- |
| [GO:0005622](http://amigo.geneontology.org/amigo/term/GO:0005622) | intracellular anatomical structure | 12 |
| [GO:0071356](http://amigo.geneontology.org/amigo/term/GO:0071356) | cellular response to tumor necrosis factor | 11 |
| [GO:0071347](http://amigo.geneontology.org/amigo/term/GO:0071347) | cellular response to interleukin-1 | 8 |
| [GO:0000977](http://amigo.geneontology.org/amigo/term/GO:0000977) | RNA polymerase II transcription regulatory region sequence-specific DNA binding | 8 |
| [GO:0043525](http://amigo.geneontology.org/amigo/term/GO:0043525) | positive regulation of neuron apoptotic process | 7 |
| [GO:0016607](http://amigo.geneontology.org/amigo/term/GO:0016607) | nuclear speck | 7 |
| [GO:0003713](http://amigo.geneontology.org/amigo/term/GO:0003713) | transcription coactivator activity | 6 |
| [GO:1900745](http://amigo.geneontology.org/amigo/term/GO:1900745) | positive regulation of p38MAPK cascade | 5 |
| [GO:1901653](http://amigo.geneontology.org/amigo/term/GO:1901653) | cellular response to peptide | 5 |
| [GO:0000785](http://amigo.geneontology.org/amigo/term/GO:0000785) | chromatin | 4 |
| [GO:0071837](http://amigo.geneontology.org/amigo/term/GO:0071837) | HMG box domain binding | 4 |
| [GO:0036464](http://amigo.geneontology.org/amigo/term/GO:0036464) | cytoplasmic ribonucleoprotein granule | 3 |
| [GO:0036488](http://amigo.geneontology.org/amigo/term/GO:0036488) | CHOP-C/EBP complex | 2 |
| [GO:1990617](http://amigo.geneontology.org/amigo/term/GO:1990617) | CHOP-ATF4 complex | 2 |
| [GO:1990037](http://amigo.geneontology.org/amigo/term/GO:1990037) | Lewy body core | 2 |
| [GO:0030014](http://amigo.geneontology.org/amigo/term/GO:0030014) | CCR4-NOT complex | 2 |
| [GO:0000307](http://amigo.geneontology.org/amigo/term/GO:0000307) | cyclin-dependent protein kinase holoenzyme complex | 2 |
