## Supplementary material for "Single-cell transcriptome sequencing for opening the blood-brain barrier through specific mode electroacupuncture stimulation": Supplementary File 19.docx (from another version of this preprint: version 1, 2025-08-03)

**Supplementary File 19. KEGG analysis for MG_cluster0 top genes only (counts top 20)**

| **Pathway_ID** | **Pathway_Name** | **S** |
| --- | --- | --- |
| [rno05202](https://www.kegg.jp/entry/rno05202) | Transcriptional misregulation in cancer | 12 |
| [rno04068](https://www.kegg.jp/entry/rno04068) | FoxO signaling pathway | 8 |
| [rno05210](https://www.kegg.jp/entry/rno05210) | Colorectal cancer | 7 |
| [rno05323](https://www.kegg.jp/entry/rno05323) | Rheumatoid arthritis | 6 |
