## Supplementary material for "Single-cell transcriptome sequencing for opening the blood-brain barrier through specific mode electroacupuncture stimulation": Supplementary File 21.docx (from another version of this preprint: version 1, 2025-08-03)

**Supplementary File 21. GO analysis for MG_cluster2 top genes only (S≥2)**

| **GO_ID** | **GO_Term** | **S** |
| --- | --- | --- |
| [GO:0005576](http://amigo.geneontology.org/amigo/term/GO:0005576) | extracellular region | 6 |
| [GO:0071310](http://amigo.geneontology.org/amigo/term/GO:0071310) | cellular response to organic substance | 4 |
| [GO:0070098](http://amigo.geneontology.org/amigo/term/GO:0070098) | chemokine-mediated signaling pathway | 4 |
| [GO:0035994](http://amigo.geneontology.org/amigo/term/GO:0035994) | response to muscle stretch | 4 |
| [GO:0045647](http://amigo.geneontology.org/amigo/term/GO:0045647) | negative regulation of erythrocyte differentiation | 3 |
| [GO:0043922](http://amigo.geneontology.org/amigo/term/GO:0043922) | negative regulation by host of viral transcription | 3 |
| [GO:0071889](http://amigo.geneontology.org/amigo/term/GO:0071889) | 14-3-3 protein binding | 2 |
| [GO:0031086](http://amigo.geneontology.org/amigo/term/GO:0031086) | nuclear-transcribed mRNA catabolic process, deadenylation-independent decay | 2 |
| [GO:0017091](http://amigo.geneontology.org/amigo/term/GO:0017091) | mRNA 3'-UTR AU-rich region binding | 2 |
| [GO:0017017](http://amigo.geneontology.org/amigo/term/GO:0017017) | MAP kinase tyrosine/serine/threonine phosphatase activity | 2 |
| [GO:0001965](http://amigo.geneontology.org/amigo/term/GO:0001965) | G-protein alpha-subunit binding | 2 |
