## Supplementary material for "Single-cell transcriptome sequencing for opening the blood-brain barrier through specific mode electroacupuncture stimulation": Supplementary File 22.docx (from another version of this preprint: version 1, 2025-08-03)

**Supplementary File 22. KEGG analysis for MG_cluster2 top genes only (counts top 20)**

| **Pathway_ID** | **Pathway_Name** | **S** |
| --- | --- | --- |
| [rno04142](https://www.kegg.jp/entry/rno04142) | Lysosome | 5 |
| [rno04062](https://www.kegg.jp/entry/rno04062) | Chemokine signaling pathway | 4 |
| [rno04137](https://www.kegg.jp/entry/rno04137) | Mitophagy - animal | 3 |
