## Supplementary material for "Single-cell transcriptome sequencing for opening the blood-brain barrier through specific mode electroacupuncture stimulation": Supplementary File 24.docx (from another version of this preprint: version 1, 2025-08-03)

**Supplementary File 24. GO analysis for MG_cluster3 top genes only (S≥2)**

| **GO_ID** | **GO_Term** | **S** |
| --- | --- | --- |
| [GO:0007568](http://amigo.geneontology.org/amigo/term/GO:0007568) | obsolete aging | 8 |
| [GO:0043025](http://amigo.geneontology.org/amigo/term/GO:0043025) | neuronal cell body | 7 |
| [GO:0001666](http://amigo.geneontology.org/amigo/term/GO:0001666) | response to hypoxia | 6 |
| [GO:1901216](http://amigo.geneontology.org/amigo/term/GO:1901216) | obsolete positive regulation of neuron death | 4 |
| [GO:0032570](http://amigo.geneontology.org/amigo/term/GO:0032570) | response to progesterone | 4 |
| [GO:0005788](http://amigo.geneontology.org/amigo/term/GO:0005788) | endoplasmic reticulum lumen | 4 |
| [GO:0019911](http://amigo.geneontology.org/amigo/term/GO:0019911) | structural constituent of myelin sheath | 4 |
| [GO:0031115](http://amigo.geneontology.org/amigo/term/GO:0031115) | negative regulation of microtubule polymerization | 3 |
| [GO:0048260](http://amigo.geneontology.org/amigo/term/GO:0048260) | positive regulation of receptor-mediated endocytosis | 3 |
| [GO:0005790](http://amigo.geneontology.org/amigo/term/GO:0005790) | smooth endoplasmic reticulum | 3 |
| [GO:0071944](http://amigo.geneontology.org/amigo/term/GO:0071944) | cell periphery | 3 |
| [GO:0051787](http://amigo.geneontology.org/amigo/term/GO:0051787) | misfolded protein binding | 3 |
| [GO:1905907](http://amigo.geneontology.org/amigo/term/GO:1905907) | negative regulation of amyloid fibril formation | 2 |
| [GO:1902949](http://amigo.geneontology.org/amigo/term/GO:1902949) | positive regulation of tau-protein kinase activity | 2 |
| [GO:0033269](http://amigo.geneontology.org/amigo/term/GO:0033269) | internode region of axon | 2 |
| [GO:0034663](http://amigo.geneontology.org/amigo/term/GO:0034663) | endoplasmic reticulum chaperone complex | 2 |
| [GO:0005528](http://amigo.geneontology.org/amigo/term/GO:0005528) | FK506 binding | 2 |
| [GO:0050750](http://amigo.geneontology.org/amigo/term/GO:0050750) | low-density lipoprotein particle receptor binding | 2 |
| [GO:0042623](http://amigo.geneontology.org/amigo/term/GO:0042623) | ATP hydrolysis activity | 2 |
