## Supplementary material for "Single-cell transcriptome sequencing for opening the blood-brain barrier through specific mode electroacupuncture stimulation": Supplementary File 25.docx (from another version of this preprint: version 1, 2025-08-03)

**Supplementary File 25. GO analysis for MG_cluster5 top genes only (S≥2)**

| **GO_ID** | **GO_Term** | **S** |
| --- | --- | --- |
| [GO:0016020](http://amigo.geneontology.org/amigo/term/GO:0016020) | membrane | 25 |
| [GO:0051607](http://amigo.geneontology.org/amigo/term/GO:0051607) | defense response to virus | 19 |
| [GO:0009615](http://amigo.geneontology.org/amigo/term/GO:0009615) | response to virus | 15 |
| [GO:0045087](http://amigo.geneontology.org/amigo/term/GO:0045087) | innate immune response | 15 |
| [GO:0045071](http://amigo.geneontology.org/amigo/term/GO:0045071) | negative regulation of viral genome replication | 10 |
| [GO:0006955](http://amigo.geneontology.org/amigo/term/GO:0006955) | immune response | 10 |
| [GO:0035458](http://amigo.geneontology.org/amigo/term/GO:0035458) | cellular response to interferon-beta | 8 |
| [GO:0042612](http://amigo.geneontology.org/amigo/term/GO:0042612) | MHC class I protein complex | 8 |
| [GO:0042605](http://amigo.geneontology.org/amigo/term/GO:0042605) | peptide antigen binding | 8 |
| [GO:0005102](http://amigo.geneontology.org/amigo/term/GO:0005102) | signaling receptor binding | 8 |
| [GO:0002474](http://amigo.geneontology.org/amigo/term/GO:0002474) | antigen processing and presentation of peptide antigen via MHC class I | 7 |
| [GO:0005789](http://amigo.geneontology.org/amigo/term/GO:0005789) | endoplasmic reticulum membrane | 7 |
| [GO:0003725](http://amigo.geneontology.org/amigo/term/GO:0003725) | double-stranded RNA binding | 7 |
| [GO:0003924](http://amigo.geneontology.org/amigo/term/GO:0003924) | GTPase activity | 7 |
| [GO:0005525](http://amigo.geneontology.org/amigo/term/GO:0005525) | GTP binding | 7 |
| [GO:0006952](http://amigo.geneontology.org/amigo/term/GO:0006952) | defense response | 6 |
| [GO:0071346](http://amigo.geneontology.org/amigo/term/GO:0071346) | cellular response to type II interferon | 5 |
| [GO:0060700](http://amigo.geneontology.org/amigo/term/GO:0060700) | regulation of ribonuclease activity | 4 |
| [GO:0034340](http://amigo.geneontology.org/amigo/term/GO:0034340) | response to type I interferon | 4 |
| [GO:0001916](http://amigo.geneontology.org/amigo/term/GO:0001916) | positive regulation of T cell mediated cytotoxicity | 4 |
| [GO:0002230](http://amigo.geneontology.org/amigo/term/GO:0002230) | positive regulation of defense response to virus by host | 4 |
| [GO:0001730](http://amigo.geneontology.org/amigo/term/GO:0001730) | 2'-5'-oligoadenylate synthetase activity | 4 |
| [GO:0044389](http://amigo.geneontology.org/amigo/term/GO:0044389) | ubiquitin-like protein ligase binding | 4 |
| [GO:0035455](http://amigo.geneontology.org/amigo/term/GO:0035455) | response to interferon-alpha | 3 |
| [GO:0035456](http://amigo.geneontology.org/amigo/term/GO:0035456) | response to interferon-beta | 3 |
| [GO:0060337](http://amigo.geneontology.org/amigo/term/GO:0060337) | type I interferon-mediated signaling pathway | 3 |
| [GO:2000406](http://amigo.geneontology.org/amigo/term/GO:2000406) | positive regulation of T cell migration | 3 |
| [GO:0035457](http://amigo.geneontology.org/amigo/term/GO:0035457) | cellular response to interferon-alpha | 3 |
| [GO:2000045](http://amigo.geneontology.org/amigo/term/GO:2000045) | regulation of G1/S transition of mitotic cell cycle | 3 |
| [GO:0001891](http://amigo.geneontology.org/amigo/term/GO:0001891) | phagocytic cup | 3 |
| [GO:0000502](http://amigo.geneontology.org/amigo/term/GO:0000502) | proteasome complex | 3 |
| [GO:0072562](http://amigo.geneontology.org/amigo/term/GO:0072562) | blood microparticle | 3 |
| [GO:0008537](http://amigo.geneontology.org/amigo/term/GO:0008537) | proteasome activator complex | 2 |
| [GO:1990111](http://amigo.geneontology.org/amigo/term/GO:1990111) | spermatoproteasome complex | 2 |
| [GO:0005771](http://amigo.geneontology.org/amigo/term/GO:0005771) | multivesicular body | 2 |
| [GO:0030670](http://amigo.geneontology.org/amigo/term/GO:0030670) | phagocytic vesicle membrane | 2 |
| [GO:0005839](http://amigo.geneontology.org/amigo/term/GO:0005839) | proteasome core complex | 2 |
| [GO:0046978](http://amigo.geneontology.org/amigo/term/GO:0046978) | TAP1 binding | 2 |
| [GO:0046979](http://amigo.geneontology.org/amigo/term/GO:0046979) | TAP2 binding | 2 |
| [GO:0097677](http://amigo.geneontology.org/amigo/term/GO:0097677) | STAT family protein binding | 2 |
| [GO:0030881](http://amigo.geneontology.org/amigo/term/GO:0030881) | beta-2-microglobulin binding | 2 |
