## Supplementary material for "Single-cell transcriptome sequencing for opening the blood-brain barrier through specific mode electroacupuncture stimulation": Supplementary File 27.docx (from another version of this preprint: version 1, 2025-08-03)

**Supplementary File 27. KEGG analysis for MG_cluster5 top genes only (20 smallest P values)**

| **Pathway_ID** | **Pathway_Name** | **S** |
| --- | --- | --- |
| [rno05168](https://www.kegg.jp/entry/rno05168) | Herpes simplex virus 1 infection | 19 |
| [rno05165](https://www.kegg.jp/entry/rno05165) | Human papillomavirus infection | 15 |
| [rno04145](https://www.kegg.jp/entry/rno04145) | Phagosome | 13 |
| [rno05160](https://www.kegg.jp/entry/rno05160) | Hepatitis C | 10 |
| [rno05170](https://www.kegg.jp/entry/rno05170) | Human immunodeficiency virus 1 infection | 10 |
| [rno05416](https://www.kegg.jp/entry/rno05416) | Viral myocarditis | 9 |
| [rno04514](https://www.kegg.jp/entry/rno04514) | Cell adhesion molecules | 8 |
| [rno05330](https://www.kegg.jp/entry/rno05330) | Allograft rejection | 7 |
| [rno05332](https://www.kegg.jp/entry/rno05332) | Graft-versus-host disease | 7 |
| [rno05320](https://www.kegg.jp/entry/rno05320) | Autoimmune thyroid disease | 7 |
| [rno04940](https://www.kegg.jp/entry/rno04940) | Type I diabetes mellitus | 7 |
