## Supplementary material for "Single-cell transcriptome sequencing for opening the blood-brain barrier through specific mode electroacupuncture stimulation": Supplementary File 28.docx (from another version of this preprint: version 1, 2025-08-03)

**Supplementary File 28. GO analysis for MG_cluster6 top genes only (S≥2)**

| **GO_ID** | **GO_Term** | **S** |
| --- | --- | --- |
| [GO:0005524](http://amigo.geneontology.org/amigo/term/GO:0005524) | ATP binding | 14 |
| [GO:0006355](http://amigo.geneontology.org/amigo/term/GO:0006355) | regulation of DNA-templated transcription | 13 |
| [GO:0006351](http://amigo.geneontology.org/amigo/term/GO:0006351) | DNA-templated transcription | 12 |
| [GO:0005730](http://amigo.geneontology.org/amigo/term/GO:0005730) | nucleolus | 10 |
| [GO:0042803](http://amigo.geneontology.org/amigo/term/GO:0042803) | protein homodimerization activity | 10 |
| [GO:0051087](http://amigo.geneontology.org/amigo/term/GO:0051087) | protein-folding chaperone binding | 9 |
| [GO:0006986](http://amigo.geneontology.org/amigo/term/GO:0006986) | response to unfolded protein | 8 |
| [GO:0006915](http://amigo.geneontology.org/amigo/term/GO:0006915) | apoptotic process | 8 |
| [GO:0010628](http://amigo.geneontology.org/amigo/term/GO:0010628) | positive regulation of gene expression | 7 |
| [GO:0051085](http://amigo.geneontology.org/amigo/term/GO:0051085) | chaperone cofactor-dependent protein refolding | 6 |
| [GO:0034605](http://amigo.geneontology.org/amigo/term/GO:0034605) | cellular response to heat | 6 |
| [GO:0014070](http://amigo.geneontology.org/amigo/term/GO:0014070) | response to organic cyclic compound | 6 |
| [GO:0044183](http://amigo.geneontology.org/amigo/term/GO:0044183) | protein folding chaperone | 6 |
| [GO:0005874](http://amigo.geneontology.org/amigo/term/GO:0005874) | microtubule | 5 |
| [GO:0005759](http://amigo.geneontology.org/amigo/term/GO:0005759) | mitochondrial matrix | 4 |
| [GO:0001671](http://amigo.geneontology.org/amigo/term/GO:0001671) | ATPase activator activity | 4 |
| [GO:0016887](http://amigo.geneontology.org/amigo/term/GO:0016887) | ATP hydrolysis activity | 4 |
| [GO:0042470](http://amigo.geneontology.org/amigo/term/GO:0042470) | melanosome | 3 |
| [GO:0035259](http://amigo.geneontology.org/amigo/term/GO:0035259) | nuclear glucocorticoid receptor binding | 3 |
| [GO:0099524](http://amigo.geneontology.org/amigo/term/GO:0099524) | postsynaptic cytosol | 2 |
| [GO:0008180](http://amigo.geneontology.org/amigo/term/GO:0008180) | COP9 signalosome | 2 |
