## Supplementary material for "Single-cell transcriptome sequencing for opening the blood-brain barrier through specific mode electroacupuncture stimulation": Supplementary File 29.docx (from another version of this preprint: version 1, 2025-08-03)

**Supplementary File 29. KEGG analysis for MG_cluster3 top genes only (counts top 20)**

| **Pathway_ID** | **Pathway_Name** | **S** |
| --- | --- | --- |
| [rno04921](https://www.kegg.jp/entry/rno04921) | Oxytocin signaling pathway | 4 |
| [rno05020](https://www.kegg.jp/entry/rno05020) | Prion disease | 3 |
| [rno05030](https://www.kegg.jp/entry/rno05030) | Cocaine addiction | 3 |
| [rno04918](https://www.kegg.jp/entry/rno04918) | Thyroid hormone synthesis | 3 |
