## Supplementary material for "Single-cell transcriptome sequencing for opening the blood-brain barrier through specific mode electroacupuncture stimulation": Supplementary File 30.docx (from another version of this preprint: version 1, 2025-08-03)

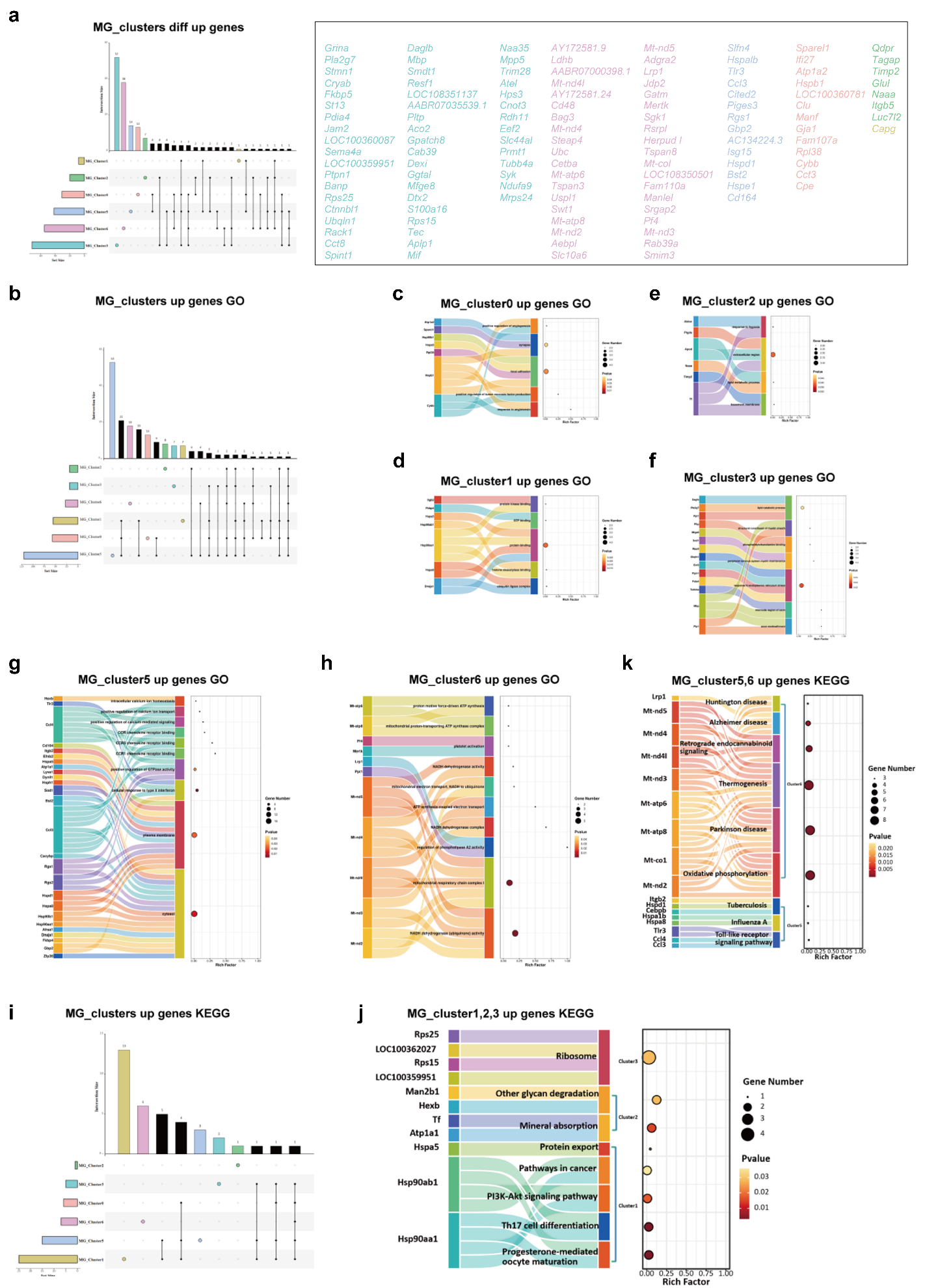

**Supplementary File 30.** **Functional enrichment analysis of unique upregulated genes in MG_clusters after electroacupuncture, related to Figure 6.** **(A)** Identical and different up-regulated genes in six subpopulations of microglia. The histograms of different colors represent different clusters and gene numbers. Coloured circles represent genes that are only upregulated in the corresponding clusters, and these genes are listed in their respective colors. **(B)** and **(I)** Crossover and separation of upregulated genes GO and KEGG in six clusters of microglia. **(C–H)** Unique important GO entries and their genes in the six clusters are displayed. **(J)** and **(K)** unique key KEGG pathways and their genes in the five clusters (MG_Cluster0 has no unique enrichment pathway). GO, Gene Ontology; KEGG, Kyoto Encyclopedia of Genes and Genomes
