## Supplementary material for "Single-cell transcriptome sequencing for opening the blood-brain barrier through specific mode electroacupuncture stimulation": Supplementary File 31.docx (from another version of this preprint: version 1, 2025-08-03)

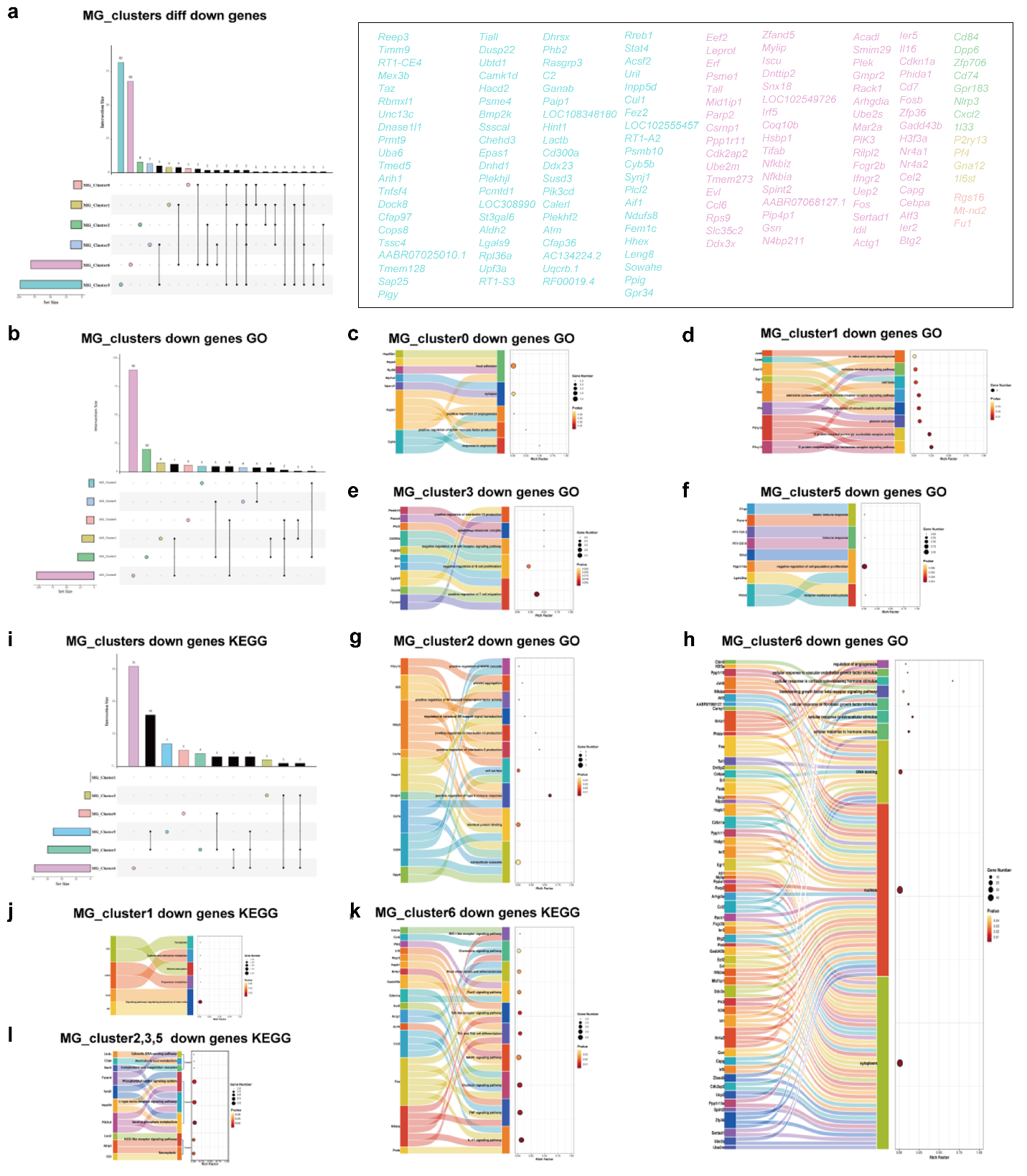

**Supplementary File 31.** **Functional enrichment analysis of unique downregulated genes in MG_clusters after EA, related to Figure 6.** **(A)** Crossover and separation of downregulated genes in six clusters of microglia. The histograms with different colors represent different clusters and gene numbers. Coloured circles represent genes that are only downregulated in the corresponding clusters, and these genes are listed in their respective colors. **(B)** and **(I)** Same and different GO and KEGG for down-regulated gene enrichment in six clusters of microglia. **(C–H)** Unique key GO entries and their genes in the six clusters are displayed. **(J–L)** Unique key KEGG pathways and their genes in the five clusters (MG_Cluster0 has no unique enrichment pathway). GO, Gene Ontology; KEGG, Kyoto Encyclopedia of Genes and Genomes
