## Supplementary material for "Single-cell transcriptome sequencing for opening the blood-brain barrier through specific mode electroacupuncture stimulation": Supplementary File 32.docx (from another version of this preprint: version 1, 2025-08-03)

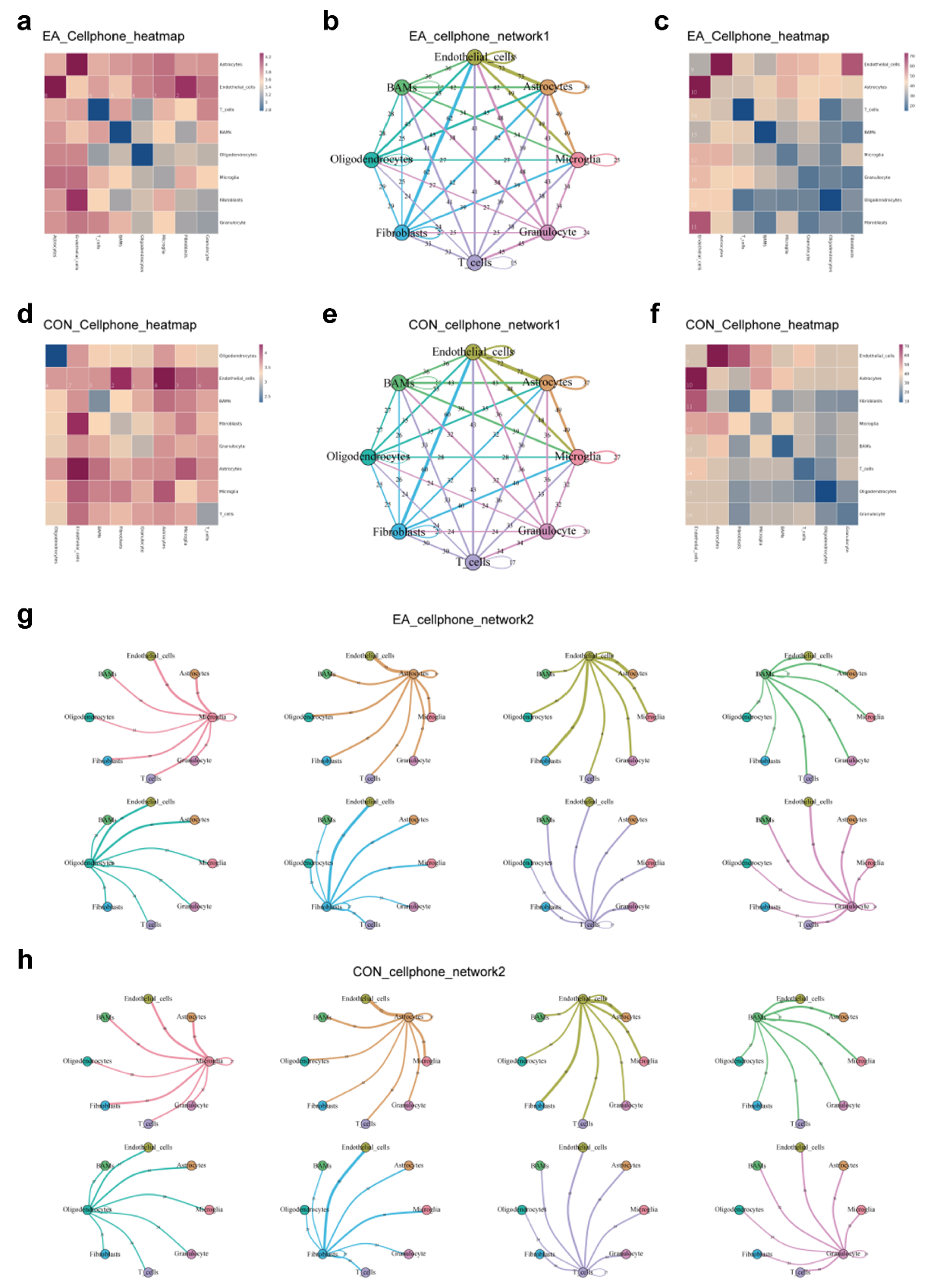

**Supplementary File 32.** **The interactions between cells, related to the section “Coordination of cell communication in the BBB following EA intervention.”** **(A–C, F)** Heatmap of the density of ligand-receptor pairs in the EA and CON groups. The redder the colour, the closer the relationship between the cell groups; **(C)** and **(B)** are the original values; **(A)** and **(F)** are the values after taking the log value. **(B, E, G, H)** Network diagram of the active degree of the ligand-receptor relationship between two cells. **(B)** and **(E)** show the overall interaction network of the two groups. **(G)** and **(H)** interactions of individual cells with other cells in both groups. The thicker the connection, the closer the correlation between the cells on the surface. CON, control; EA, electroacupuncture
