## Supplementary material for "Single-cell transcriptome sequencing for opening the blood-brain barrier through specific mode electroacupuncture stimulation": Supplementary File 33.docx (from another version of this preprint: version 1, 2025-08-03)

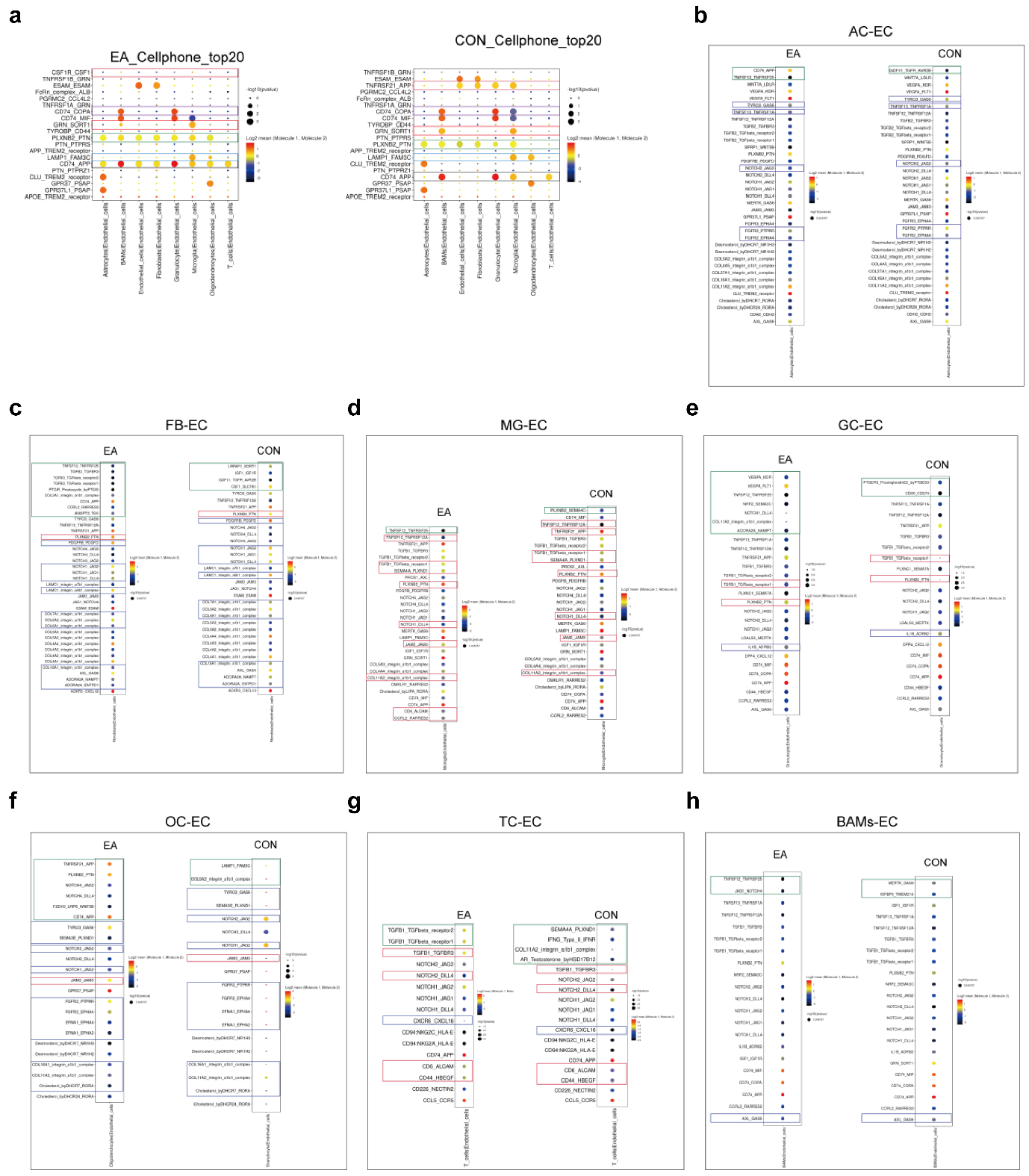

**Supplementary File 33.** **Cell communication with endothelial cells, related to the section “Coordination of cell communication in the BBB following EA intervention.” (A)** The top 20 ligand receptor pairs with mean value in the EA and CON groups, respectively. Different colored boxes are used to indicate pairs with differences between the groups. **(B–H)** Detailed ligand receptor pairs for communication between other cells to endothelial cells. The depth of the colors in the image represents the strength of the corresponding interaction, and the size of the dots represents the significance of the *P*-value. The green box represents pairs that are only present in a specific group (EA or CON), the red box represents pairs that are upregulated in the EA group (mean or *P* value), and the blue box represents pairs that are downregulated in the EA group. CON, control; EA, electroacupuncture
